# Predictive modeling of long non-coding RNA chromatin (dis-)association

**DOI:** 10.1101/2020.12.15.422063

**Authors:** Evgenia Ntini, Stefan Budach, Ulf A Vang Ørom, Annalisa Marsico

## Abstract

Long non-coding RNAs (lncRNAs) are involved in gene expression regulation in *cis* and *trans*. Although enriched in the chromatin cell fraction, to what degree this defines their broad range of functions remains unclear. In addition, the factors that contribute to lncRNA chromatin tethering, as well as the molecular basis of efficient lncRNA chromatin dissociation and its functional impact on enhancer activity and target gene expression, remain to be resolved. Here, we combine pulse-chase metabolic labeling of nascent RNA with chromatin fractionation and transient transcriptome sequencing to follow nascent RNA transcripts from their co-transcriptional state to their release into the nucleoplasm. By incorporating functional and physical characteristics in machine learning models, we find that parameters like co-transcriptional splicing contributes to efficient lncRNA chromatin dissociation. Intriguingly, lncRNAs transcribed from enhancer-like regions display reduced chromatin retention, suggesting that, in addition to splicing, lncRNA chromatin dissociation may contribute to enhancer activity and target gene expression.

**Highlights:**

- Chromatin (dis-)association of lncRNAs can be modeled using nascent RNA sequencing from pulse-chase chromatin fractionation
- Distinct physical and functional characteristics contribute to lncRNA chromatin (dis-)association
- lncRNAs transcribed from enhancers display increased degree of chromatin dissociation
- lncRNAs of distinct degrees of chromatin association display differential binding probabilities for RNA-binding proteins (RBPs)

## Introduction

Bidirectional nascent RNA transcription is a prominent characteristic of active enhancers, leading to the production of short-lived non-coding RNA transcripts termed eRNAs. eRNAs are short and non-spliced, thus unstable, potentially terminated by the Integrator complex^1^ and subjected to rapid exosome degradation^2^, thereby contributing to their observed chromatin enrichment and eliminated detection in steady-state whole-cell RNA data. eRNA production, measured by nascent RNA-sequencing, along with DNase I hypersensitivity and distinct histone marks (H3K27Ac, H3K4me1) (and CBP/p300 binding) demarcate active enhancers^2–6^.

Intriguingly, a small subset of bidirectionally transcribed enhancers, about ~3 to 5 %, produce a more stable and spliced long non-coding RNA (lncRNA) elongating in one direction^3,7^ (& Tan and Marques, biorXiv 2020), while about one third to one fourth of annotated lncRNAs overlap enhancer-like regions^8^. Those enhancer-associated lncRNAs (*elncRNAs*) are associated with stronger enhancer activity (aka. higher nascent RNA transcription, H3K27Ac histone mark, DNase accessibility) and their expression is associated with changes in putative target gene expression and local chromatin structure. This suggests that elncRNA production contributes to gene expression regulation *in cis*^3,7^. However, to what degree elncRNAs remain chromatin-associated (in a manner analogous to the observed eRNA chromatin enrichment), and the degree to which their function depends on their chromatin (dis-)association remains obscure. In addition, the mechanistic basis of their exerted regulation on target gene expression *in cis* is not well characterized, and it remains an open question whether all elncRNAs would follow the same mechanistic mode in gene expression regulation. For instance, we showed that the lncRNA *A-ROD* transcribed from an active enhancer at the anchor point of a chromosomal loop in *MCF-7* cells enhances the expression of its target gene *DKK1* upon its post-transcriptional chromatin dissociation and within a pre-established chromosomal proximity. Enforcing *A-ROD* chromatin retention, by splicing inhibiting morpholinos or targeting polyadenylation, suppresses target gene expression, suggesting that chromatin dissociation is an important feature of lncRNA mediated gene expression regulation *in cis*^9^.

A substantial portion of lncRNAs are enriched in the chromatin fraction, presumably tethered at their sites of transcription through elongating (transcriptionally engaged) Pol II, and are involved in regulation of proximal gene expression *in cis*^10–12^. However, intriguingly, lncRNAs transcribed from the anchor points of chromosomal loops and enhancer-like regions show significantly lower chromatin—to—nucleoplasmic enrichment at steady state^9^. This may indicate that the process of chromatin dissociation, which relies on (co-transcriptional) RNA maturation steps, could be important for the function of many enhancer-transcribed lncRNAs, acting *in cis* within the spatial proximity of pre-established chromosomal loops^13^.

A recent study additionally implicated U1 snRNP binding as a means of chromatin tethering for lncRNAs: lncRNA exonic sequences are enriched in U1 recognition sites, while their gene bodies are depleted from 3’ splice sites (compared to mRNAs). This leads to persistent U1 snRNP binding —due to poor or inefficient splicing efficiency—, which through additional protein interactions with transcriptionally engaged Pol II, contributes to co-transcriptional lncRNA tethering (or post-transcriptional retargeting) to chromatin^14^. Intriguingly, compared to other lncRNAs that are not enhancer-associated, elncRNAs display conserved splice sites and significantly higher splicing efficiency, which is associated with local changes in chromatin states and positively impacts their cognate enhancer activity^3,7,13^. Yet, a correlation between elncRNA splicing and chromatin-association/dissociation has not been clarified. Although recent bioinformatics approaches strongly infer an impact of elncRNA processing on enhancer activity, the role of elncRNA chromatin (dis-)association has not been systematically examined.

In this work, we have combined pulse-chase metabolic labeling with chromatin fractionation and transient transcriptome sequencing to follow nascent RNAs from the point of their transcription to their chromatin release into the nucleoplasm. We have incorporated several parameters, physical and functional characteristics, in machine learning models to predict distinct degrees of chromatin (dis-)association, and examined the relationship between lncRNA chromatin dissociation and enhancer activity. Thus, two important questions are addressed here: First, what are the parameters that contribute to distinct degrees of lncRNA chromatin association or chromatin tethering. Second, whether increased chromatin dissociation of certain lncRNAs could imply a functional potential, for instance by having an impact on— or shaping enhancer activity.

## Results

### Modeling chromatin (dis-)association of nascent RNA transcripts

To follow nascent RNA transcripts from their synthesis to their post-transcriptional chromatin dissociation we performed nascent RNA sequencing from the chromatin-associated and nucleoplasmic fraction. We performed 4-thiouridine (4-SU) metabolic labeling of MCF-7 cells for an 8 min pulse, followed by 5, 10, 15 and 20 min uridine chase (Methods). To additionally capture nascent RNA Pol II transcription in a high resolution and follow transcription dynamics, we fragmented RNA prior to isolation of nascent RNA. Thus, our approach is similar to ‘transient transcriptome sequencing’ (*TT-seq*^15^) but coupled with chromatin fractionation and pulse-chase labeling.

To model chromatin dissociation we extracted read coverage from the last exon, as we did not block new transcription initiation events during the pulse-chase experiment (in the case of overlapping transcript isoforms the longest transcript was selected; Methods). This was done to minimize transcriptional input from new transcription initiation events during the pulse-chase time period and be closer to the transcript 3’ end, thus better reflecting capturing full-length transcripts. We see chromatin-associated read coverage decrease over time and nucleoplasmic read coverage increase (Suppl. Figure S1A). We determined chromatin association as the ratio CHR/(CHR+NP) at each time point and kept only transcripts with a defined ratio 0 to 1 at all time points (NAs discarded, n = 15,157 transcripts). As expected, we see an overall decrease in the transcript chromatin association over pulse-chase time (Figure 1A). We therefore fitted these ratios on an exponential decay curve to extract a ‘chromatin-association halftime’: [halftime = - (Intercept+ln2) / k]. For further analysis, we kept only entries that fit the exponential decay curve with a p-value < 0.05 (n = 12,391 transcripts, of which 2,077 are lncRNAs; Methods). We then split the dataset in 3 equal-size quantiles based on the calculated chromatin association halftime, i.e. ‘fast’, ‘medium’ and ‘slow’ released transcripts (Figure 1B, 1C, Suppl. Figure S1B) (the latter correspond to chromatin-retained transcripts). Alternatively, transcripts were clustered into 3 groups of fast, medium and slow released using the CHR/(CHR+NP) ratios from the five time points as an input to k-means clustering (Suppl. Figure S1C). In general, there is a good agreement between the two methods of grouping, with the 3 groups of k-means clustering showing corresponding chromatin association halftimes (Suppl. Fig. S1D). Although significantly shorter and with a smaller number of exons as previously reported^16^, lncRNAs show on average greater chromatin association halftimes compared to mRNAs (Suppl. Figures S1E-G). Chromatin association halftimes extracted this way reflect the chromatin association ratios at steady state (Suppl. Figure S1H). We find 872 fast, 499 medium and 706 slow-released lncRNAs (Suppl. Table 1). Two representative lncRNAs are *A-ROD* as a fast-released, and *PVT1* as a slow-released, chromatin-retained transcript (Figure 1D).

**Figure 1.**
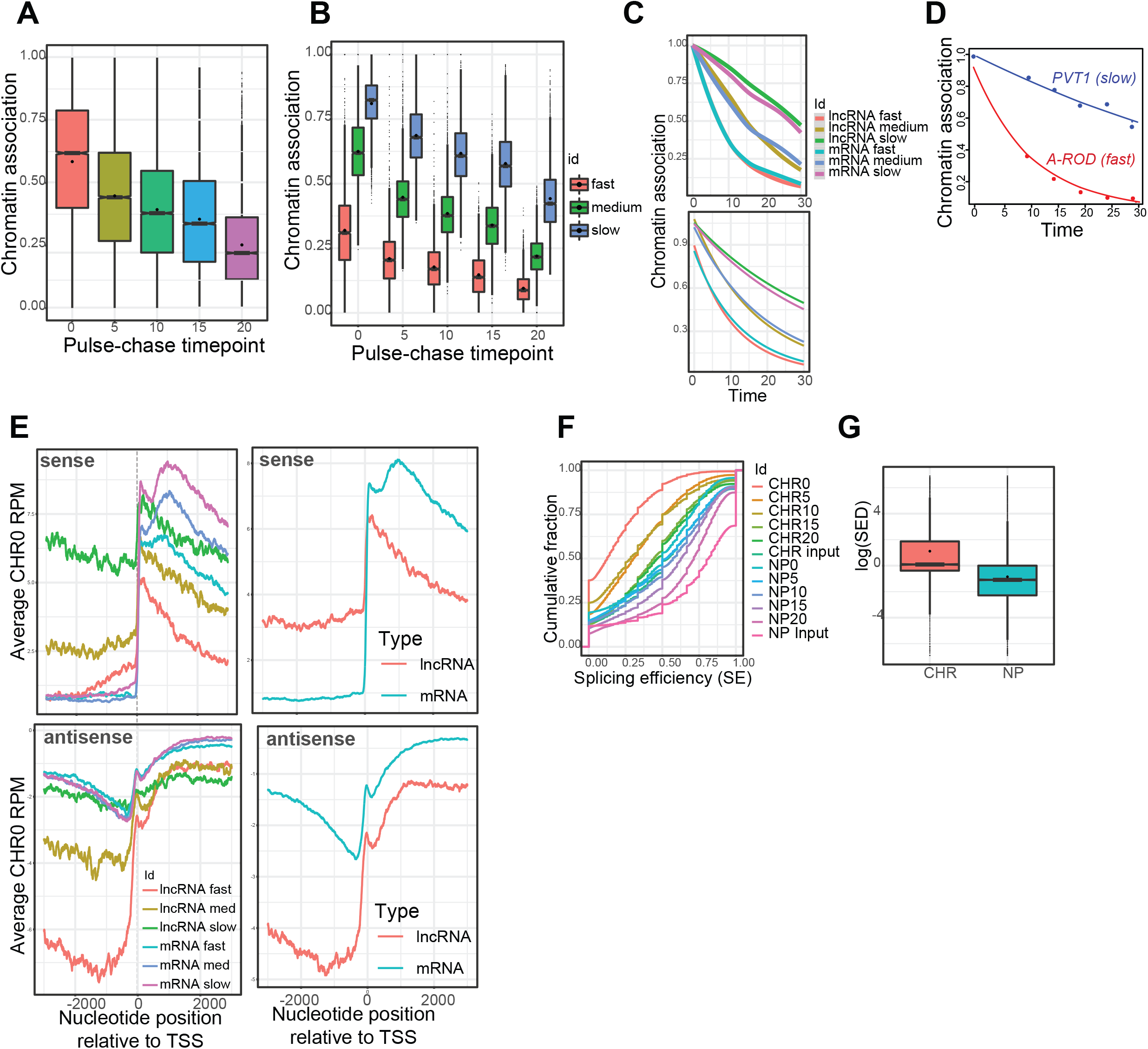
Measuring chromatin association of nascent RNA transcripts. (A) Distribution of chromatin association ratios at different pulse-chase time points. (B) Same as in (A) but split for fast, medium and slow-released transcripts. (C) Loess curve of chromatin association drawn based on the raw ratios (upper panel) and after fit on an exponential decay (lower panel). (D) Exponential decay fit of the chromatin association over time for two representative lncRNAs, A-ROD (fast-/efficiently released) and PVT1 (slow-released/chromatin retained). (E) Metagene analysis of ‘CHR0’ strand-specific read coverage (chromatin-associated nascent RNA sequencing from time point zero) in a ±3 Kb window around TSS. (F) Cumulative distribution function (CDF) curves of intron splicing efficiencies measured in all analyzed samples. (G) Distribution boxplots of intron splicing efficiency dynamics (SED) measured in the chromatin-associated (CHR20-CHR0) and nucleoplasmic fraction (NP20-NP0). Co-transcriptional SED is significantly higher compared to post-transcriptional SED (p-value < 2.2e-16).

Nascent RNA sequencing from the chromatin associated fraction allows to follow Pol II transcriptional dynamics in high resolution: application of a short metabolic pulse and RNA fragmentation prior nascent RNA purification, as in the original TT-seq protocol^15^, combined with chromatin fractionation further enriches for nascent RNA reads^17^. By metagene analysis to profile nascent RNA transcription, we obtain Pol II transcriptional profiles similar to the original TT-seq^15^ (Figure 1E). Nascent RNA sequencing from the chromatin-associated fraction also captures promoter-associated divergent transcription producing short unstable antisense transcripts (PROMPTs)^18,19^. We note here that lncRNA loci produce higher upstream antisense transcription compared to mRNAs which extends beyond the typical PROMPT length (~200 nt) (Figure 1E, lower right panel). This is most probably because many lncRNAs arise upstream and antisense to protein coding genes (and the observed upstream antisense signal is due to the associated mRNA transcription). Interestingly, we observe that fast-released lncRNAs display stronger upstream antisense signal, suggesting that fast-released lncRNAs originate more often upstream antisense of protein coding genes. Indeed, by plotting the interdistance to closest antisense protein coding gene TSS, we find that fast-released lncRNAs display on average significantly smaller values (Supplementary Figure S1 I). Notably, about half of fast-released lncRNAs originate within less than 1 kb antisense to mRNA TSS (either upstream or internal antisense) (Supplementary Figure S1 J). An example is the fast-released lncRNA *GATA3-AS1* transcribed upstream and antisense of *GATA3* (Supplementary Figure S1 M). As expected, ENCODE annotated lncRNAs with the biotype ‘antisense’ are enriched in fast-released transcripts (odds ratio 1.4657, p-value = 4.217e-06), whereas *de novo* assembled lncRNA transcripts from the chromatin-associated data not overlapping ENCODE annotations (Methods) are enriched in the slow released/chromatin-retained transcripts (odds ratio 2.070958, p-value 1.917e-11).

### Nascent RNA sequencing coupled with chromatin fractionation reveals major co-transcriptional nascent RNA processing and some degree of post-transcriptional splicing

Nascent RNA sequencing from the chromatin-associated and nucleoplasmic fraction at different pulse-chase time points allows to track the progress of co- and post-transcriptional splicing. To measure splicing we used high confidence introns (Methods) and extracted splicing efficiency by calculating the ratio of split to non-split reads at the 3’ splice site as in ref^20^. By plotting the cumulative fraction of intron splicing efficiencies from all time points and samples, we observe that most of the introns undergo extensive splicing co-transcriptionally while at chromatin, within the first 10-15 min of transcription (Figure 1F, Suppl. Figure S1K), and co-transcriptional splicing efficiency dynamics^21^ (SED; Methods) is significantly higher compared to post-transcriptional nucleoplasmic SED (Figure 1G). These results are in agreement with recent reports that the majority of splicing occurs co-transcriptionally (Reimer et al., bioRxiv 2020). We then calculated the extent of post-transcriptional splicing (after chromatin dissociation) relative to co-transcriptional splicing (while at chromatin) (as the difference between chromatin and nucleoplasmic splicing efficiency, normalized to chromatin; Methods). This was done at intron and transcript level (by extracting a mean processing efficiency from a transcript’s high-confidence introns; Methods). We observe that introns of fast-released lncRNAs, and respectively fast-released lncRNA transcripts undergo the least additional post-transcriptional splicing upon chromatin dissociation (Supplementary Figure S1 L i-ii), suggesting that most of their processing has been concluded co-transcriptionally while at chromatin. Overall, mRNAs may undergo some further post-transcriptional processing to a higher degree compared to lncRNAs (Suppl. Fig. S1 L iii). This is in agreement with recent findings using single molecule RNA FISH suggesting that some post-transcriptional splicing can occur upon chromatin dissociation, after transcription is completed, and potentially while nascent RNA transcripts localize to speckles (Coté et al., bioRxiv 2020). That slow-released transcripts show overall more extensive post-transcriptional splicing (Suppl. Fig. S1 L) is also in agreement with a model where completely synthesized nascent RNA transcripts move slowly through a transcription site proximal zone (without being tethered to chromatin or the transcription site anymore), while they can undergo additional post-transcriptional splicing (Coté et al., bioRxiv 2020).

### Different degrees of chromatin association correlate with distinct physical (and functional) characteristics of nascent RNA transcripts

We observe that lncRNAs show on average significantly lower co-transcriptional splicing efficiency compared to mRNAs (Figure 2A left), which is in agreement with what was previously reported measuring splicing using either steady-state or nascent RNA data^20,22^. In addition, fast-released mRNAs, but not lncRNAs, show on average higher mean transcript splicing efficiencies compared to slow-released/chromatin-retained transcripts (Figure 2A right). However, the minimum splicing efficiency per transcript (i.e. splicing efficiency of the worst spliced intron) is significantly higher for fast-released lncRNAs compared to chromatin-retained transcripts, suggesting that splicing of a slowly or inefficiently processed intron may act as a kinetic ‘bottleneck’ for nascent RNA transcript chromatin dissociation (Figure 2B). As expected, lncRNAs show on average significantly higher alternative splicing compared to mRNAs (intron *psi* value extracted as in ref^21^), and chromatin-retained lncRNAs undergo significantly higher alternative splicing compared to fast-released transcripts (Suppl. Fig. S2A).

**Figure 2.**
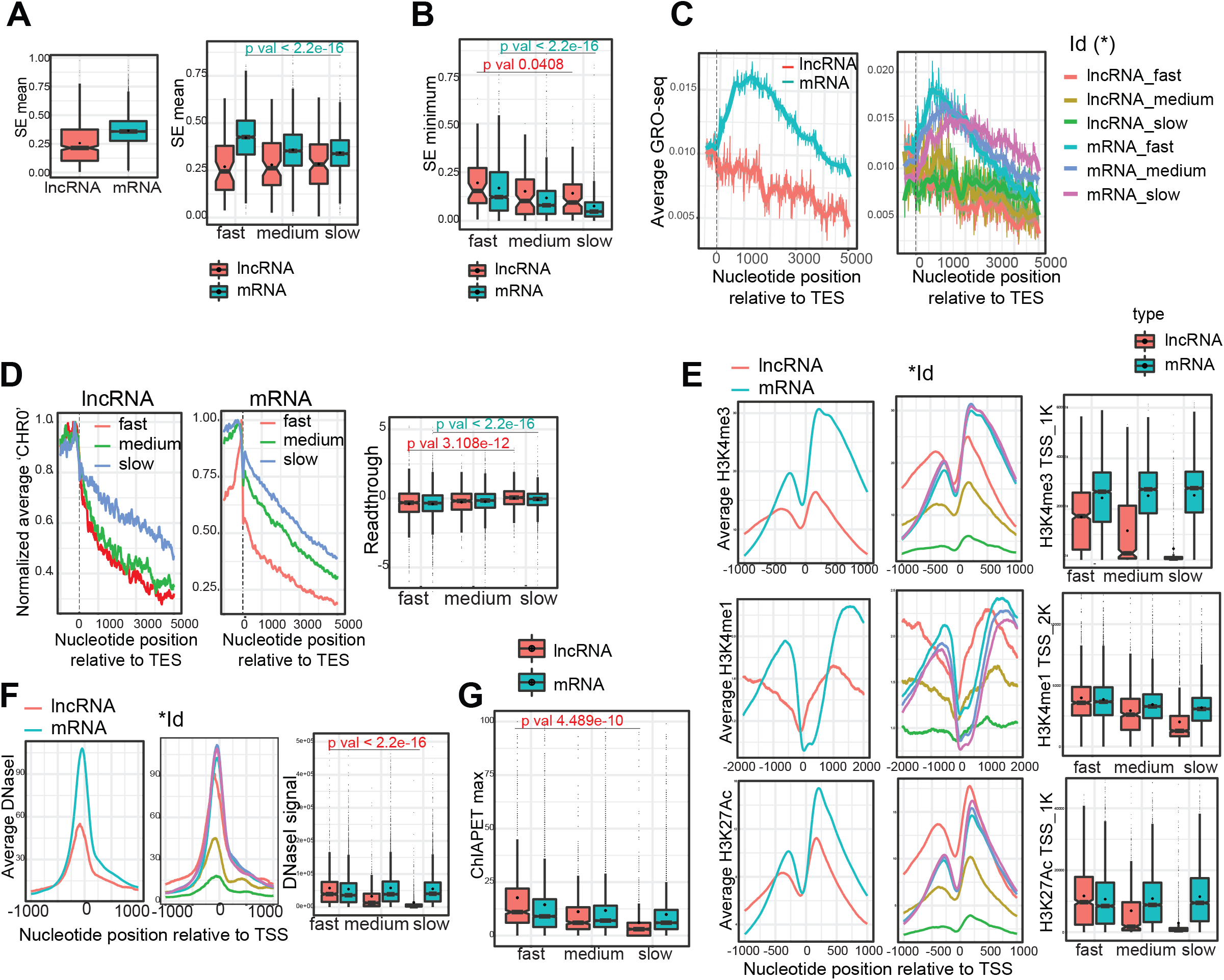
Different degrees of chromatin association correlate with distinct physical and functional characteristics of nascent RNA transcripts. (A) Transcript mean splicing efficiency. P-value < 2.2e-16 for fast-vs. slow-released mRNAs; non-significant (NS) for lncRNAs. (B) Transcript minimum splicing efficiency (i.e. splicing efficiency of worst spliced intron). P-value < 2.2e-16 fast-vs. slow-released mRNAs, p-value 0.0408 fast-vs. slow-released lncRNAs. (C) Metagene analysis of GRO-seq read coverage (only sense strand plotted) in a window −500 bp to +5 Kb around transcript end site (TES) (ContextMap extracted pA site). (D) Metagene analysis of ‘CHR0’ sense strand-specific read coverage (chromatin-associated nascent RNA sequencing from time-point zero) in the window −1 Kb to +5 Kb around TES. The average read coverage per nucleotide position is normalized to the position with the maximum read coverage within each group, and plotted separately for lncRNAs (left panel) and mRNAs (middle panel). Right panel: Boxplot distribution (plotted in log scale) of transcriptional readthrough for the different groups, measured as the ratio of ‘CHR0’ sense strand-specific read coverage 5 Kb downstream to 1 Kb upstream of TES (p-value = 3.108e-12 fast vs. slow released lncRNAs and p-value < 2.2e-16 fast vs. slow released mRNAs). (E) Metagene analysis of histone marks average profiles around the TSS of different groups of nascent RNA transcripts (left panels: split lncRNAs vs. mRNAs, middle panels: split distinct degrees of chromatin association i.e. fast/medium/slow-released lncRNAs and mRNAs). Right panels: Boxplot distribution of promoter-associated histone mark signal around TSS (p-value < 2.2e-16 fast vs. slow-released lncRNAs; NS for mRNAs). (F) Metagene analysis of DNase-seq signal around TSS (left, middle panels) and the respective boxplot distribution (right panel, p-value < 2.2e-16 fast vs. slow-released lncRNAs; NS for mRNAs). (G) Promoter-overlapping ChIA-PET maximum scores (fast vs. slow released lncRNAs p-value 4.489e-10).

By extracting the promoter-associated transcriptional pausing index (Methods) using MCF-7 available Pol II P-Ser2 ChIP-seq or GRO-seq data^23^, we find that mRNAs show significantly higher pausing index as previously reported^20,24^ (Suppl. Fig. S2B-D, S2F). Interestingly, fast-released lncRNAs, but not mRNAs, display higher pausing index compared to chromatin-retained transcripts, suggesting that transcriptional activity *per se* may relate to lncRNA chromatin dissociation or tethering (Suppl. Fig. S2C, S2D). In agreement, we find significantly different levels of transcriptionally engaged Pol II over the first Kb downstream of TSS for fast versus slow-released lncRNAs, but not mRNAs which are overall more transcriptionally active (Supplementary Fig. S2E, S2F). Taken together, these observations are in agreement with a recent report that promoters of lncRNAs show distinct transcriptional burst kinetics compared to mRNAs (lower burst frequencies; Johnsson et al., bioRxiv 2020), and suggest that within lncRNAs, promoters of fast-released transcripts tend to be more transcriptionally active and display higher degree of Pol II pausing compared to chromatin-retained lncRNA transcripts. Interestingly, transcriptional pausing index was previously associated with lncRNA nuclear export^24^. We note here that by extracting transcription bi-directionality score (or divergent-transcription score) using GRO-seq (as antisense/sense signal from 1 Kb around TSS) we reach the same conclusion by using chromatin associated nascent RNA sequencing from time point 0 (‘CHR0’, Figure 1E), showing that fast-released lncRNAs display significantly higher antisense (divergent) transcription (Supplementary Figure S2 G), which is most probably due to their enrichment in originating near and antisense of protein-coding gene TSS (Suppl. Figures S1 I-J).

Chromatin dissociation of nascent RNA transcript is coupled to transcription termination and 3’ end formation. We thus generated transcription metagene profiles around the transcript 3’ end site (TES) using ChIP-seq signal from transcriptionally engaged Pol II phosphorylated at Ser2 (P-ser2 Pol II occupancy) or strand-specific GRO-seq read coverage. To account for annotation discrepancies, we extracted *de novo* putative (pA) polyadenylation sites from ENCODE available MCF-7 nuclear polyA+ RNA-seq data using ContextMap^25^. Although in general there is good agreement between the annotated transcript 3’ ends and the *de novo* extracted pA sites (Suppl. Fig. S2 H), for increased positional accuracy we used the latter for further analyses (i.e. assigned a transcript 3’ end to closest and stronger ContextMap predicted pA site, Methods). P-Ser2 Pol II metagene profiles around TES resemble the ones obtained by mNET-seq^20^, revealing polyadenylation associated Pol II pausing in a 2 Kb window downstream of TES of mRNAs, but not lncRNAs (Supplementary Figure S2 I, left). In conjunction, mRNAs display significantly higher transcription termination index compared to lncRNAs, as previously reported^20^ (Supplementary Figure S2 I, right). GRO-seq metagene analysis profiles of transcriptionally engaged Pol II verify these results (Figure 2C, Suppl. Figure S2 K). In particular, we find no significant difference in the transcription termination index (extracted using GRO-seq) between fast and slow released mRNA transcripts, indicating no significant differences in polyadenylation-associated TES-downstream Pol II pausing (Suppl. Figure S2 K). This could suggest no significant differences in transcription termination efficiencies *per se*. Yet, by extracting a Pol II ‘travel index’ (as the ratio of strand-specific GRO-seq signal from the region 2.5 to 5 Kb downstream of TES to the first 2.5 Kb downstream of TES where the polyadenylation-associated pausing resides^20^; Methods), we note that Pol II of slow-released transcripts tends to travel further beyond the polyadenylation-associated pausing site, which would be in support of chromatin tethering via ongoing transcription^10^ (Suppl. Figure S2 L; S2 J; Figure 2C right panel) (or that ongoing transcription may contribute to chromatin tethering and slow release of nascent RNA transcript). In the case of lncRNAs, we do not observe a polyadenylation-associated TES-downstream Pol II accumulation or pausing, which is in agreement with mNET-seq data suggesting polyadenylation-independent transcription termination modes^20^. Notably, and more evidently observed in normalized metagene transcriptional profiles, Pol II tends to transcribe further beyond the TES of slow-released lncRNA transcripts (Suppl. Figure S2 J, lower panels). In agreement, by extracting travel (readthrough) ratios using chromatin-associated nascent RNA-seq from time point 0 (‘CHR0’) we find that slow-released chromatin-retained nascent RNA transcripts, either mRNAs or lncRNAs, exhibit higher readthrough transcription (Figure 2D). Taken together with the observed inefficient splicing of slow-released transcripts (Figure 2A-B), these results are in agreement with a crosstalk between splicing, transcription and transcription termination^26,27^, and with recent findings that inefficient splicing associates with readthrough transcription (Reimer et al., biorXiv 2020).

### Different degrees of chromatin association demarcated by distinct chromatin states

We then examined whether distinct degrees of nascent RNA transcript chromatin association would relate to distinct chromatin states. Notably, for all histone marks associated with transcriptional activity (H3K4me3, H3K4me1, H3K27Ac) we see significant differences in the promoter regions around the TSS of fast, medium and slow-released lncRNAs, but not for mRNAs (Figure 2E). By extracting the ratio H3K4me1 to H3K4me3 around the TSS, we observe that the fast-released lncRNAs resemble mRNAs in terms of promoter activity (Suppl. Fig. S3 A), while slow-released/chromatin-retained lncRNAs display on average higher signals of repressive histone marks like H3K9me3 and H3K27me3 (Suppl. Fig. S3 B). Profiles of total Pol II occupancy (POL2RA ChIP-seq) confirm the differences in the transcriptional activity among distinct degrees of chromatin association for lncRNAs (Suppl. Figure S3 C). Notably, fast-released lncRNAs are transcribed from regions with significantly greater chromatin accessibility (measured by DNase-seq, Figure 2F), and display significantly higher CTCF and YY1 binding (for the latter, Avocado calculated binding probability^28^; Methods) (Suppl. Figure S3 D-E). This is important, as both factors are associated with chromatin looping, and YY1 in particular promotes enhancer-promoter chromatin loops by forming protein dimers and facilitating DNA interactions^29^. The negative correlation between the extracted chromatin association halftime and looping scores (i.e. promoter-overlapping ChIA-PET nodes; Methods) is greater for lncRNAs compared to mRNAs (Pearson’s correlation - 0.267 vs. −0.107, respectively). Notably, promoters of fast-released lncRNAs display significantly higher ChIA-PET scores (Figure 2G, Suppl. Fig. S3 F), indicating that they are/tend to be transcribed from the anchor points of chromosomal loops.

### Enhancer-associated lncRNAs (elncRNAs) do not remain chromatin-associated

We therefore examined the association of distinct degrees of lncRNA chromatin dissociation with enhancer activity. For this purpose we used the FANTOM5^2,6^ human ‘permissive’ enhancers expanded by transcribed enhancers defined by NET-CAGE^4^. We filtered that these enhancers should be transcriptionally active in MCF-7 cells by GRO-seq measurement, ending up with 10,008 high-confidence bidirectionally transcribed enhancers (Fig. 3A). About 2.5 % of bidirectionally transcribed enhancers have an lncRNA TSS (derived from the analyzed dataset 2,077 lncRNAs) within an interdistance < 2 kb which is reminiscent to what was previously reported^3,7^. Thus, those lncRNAs can be regarded as enhancer-associated elncRNAs^7^ and their cognate enhancers as la-EPCs^3^. Notably, fast-released lncRNAs are significantly enriched in elncRNAs (odds ratio 1.68, p-value 0.0001398). On the other hand, ~7.6 % of the bidirectionally transcribed enhancers have an mRNA TSS (from the 10,314 analyzed) within less than 2 Kb interdistance, however fast-released mRNAs are not enriched in this subset (odds ratio 0.97). This suggests that transcribed enhancers are more likely to be associated with a fast-released lncRNA. In other words, when bidirectionally transcribed enhancers are associated with an lncRNA (at ~3-5 %), then this is more likely to be a fast-released lncRNA transcript. Analogously, we find that elncRNAs (defined at an interdistance < 2 kb to closest enhancer midpoint; Fig. 3C) are enriched in fast-released lncRNAs (odds ratio ~1.8, p-value 0.002586), whereas mRNAs with an interdistance < 2 kb to closest enhancer midpoint are not enriched in fast-released mRNAs (Fig 3B). Notably, elncRNAs show significantly higher association to anchor points of chromatin loops (measured by score of overlapping ChIA-PET nodes; Fig. 3D), and display on average significantly lower chromatin-association halftimes (p-value = 5.116e-06; Fig. 3E), (while, as a control, fast-released mRNAs are not enriched in interdistances less than 2 kb to enhancer midpoint: odds ratio 0.8409119, p-value 0.03862). This is similar to what was previously published, that lncRNAs transcribed from enhancer-like regions display on average higher ChIA-PET scores on their overlapping promoter regions^9^. Notably, although lncRNAs as a class display higher chromatin association halftimes compared to mRNAs, elncRNAs escape this rule by showing significantly lower chromatin association halftimes (Fig. 3E), which is in agreement with elncRNAs being enriched in fast-released transcripts. In conclusion, we show here that enhancer-associated or rather, enhancer-transcribed lncRNAs (elncRNAs, equivalent to la-EPCs^3^), in addition to increased splicing efficiencies^3,7^, also show increased degrees of chromatin dissociation.

**Figure 3.**
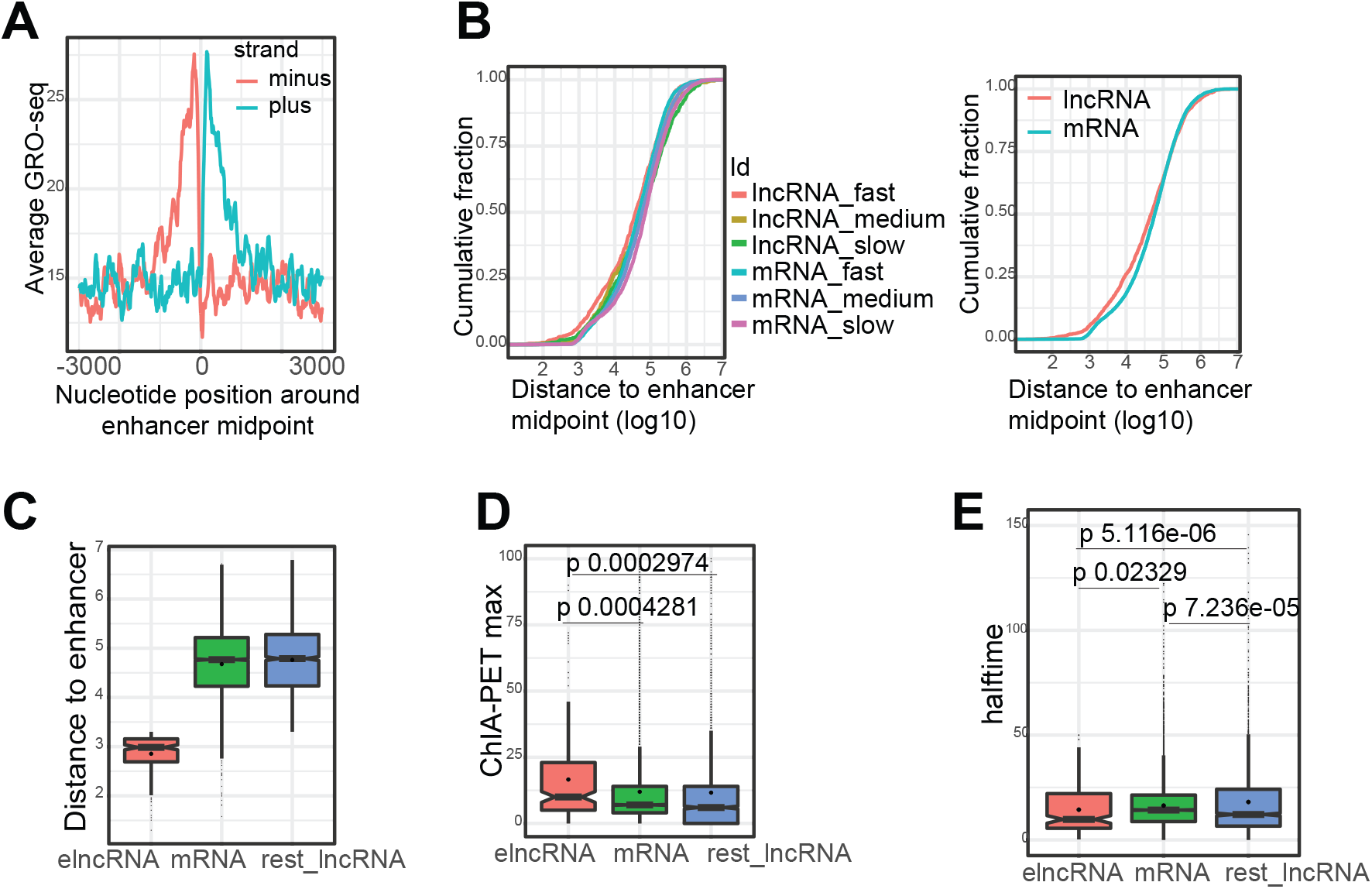
Enhancer-associated elncRNAs are enriched in fast-released transcripts. (A) Profile of nascent RNA transcription (GRO-seq) over bidirectionally transcribed enhancers in MCF-7. (B) Cumulative plots of interdistances of transcript TSS to closest enhancer midpoint. (C) Distribution of TSS interdistances (log10 bp) to closest enhancer midpoint for elncRNAs, mRNAs and lncRNAs not associated to active enhancers. (D) elncRNAs show significantly higher ChIA-PET interaction scores compared to mRNAs (p-value 0.0004281) and to lncRNAs not associated with active enhancers (p-value 0.0002974). (E) elncRNAs show significantly lower chromatin association halftimes (p-value 0.02329 to mRNAs and 5.116e-06 to rest lncRNAs; p-value 7.236e-05 rest lncRNAs to mRNAs).

### Prediction of lncRNA chromatin dissociation in machine learning models

We then incorporated several of the functional and physical characteristics in machine learning to predict chromatin dissociation of lncRNAs. We applied logistic regression with a ten-times cross-validation to predict fast versus slow-released transcripts (Figure 4A lncRNA, 4B mRNA). In agreement with the distribution of the individual parameters (Figure 2, Suppl. Fig. S2, S3) we find that the transcript exon density (previously used as a proxy for splicing activity^3^), splicing efficiency of the transcript’s worst processed intron and chromatin states associated with promoter transcriptional activity (H3K4me3 and H3K4me1) have significant coefficients in predicting fast-released lncRNAs, whereas SNRP70 enrichment across the locus (mean of fold-enrichment from ChIP-seq peaks; Methods) define slow-released, chromatin-retained lncRNAs. The latter is in agreement with Yin et al. (2020) suggesting U1-mediated chromatin retention of inefficiently processed transcripts^14^. That P-Ser2 Pol II coverage over gene body is significant in predicting slow-released lncRNAs could confirm that slow-released lncRNAs are tethered to chromatin through transcriptionally engaged Pol II^10^ and that transcriptional activity could contribute to U1 snRNP-mediated tethering of inefficiently processed transcripts^14^. In contrast to the exon density (which reflects overall splicing activity) and the splicing efficiency of the worst spliced intron, potentially acting as a kinetic bottleneck in nascent RNA transcript chromatin release, and while those two parameters confidently predict lncRNA chromatin dissociation, we (paradoxically) find the transcript’s mean splicing efficiency as an important predictive parameter of chromatin association. This could be explained if some (or one, or few) easy to process introns achieve high splicing efficiency during their prolonged stay on chromatin, thereby contributing to increasing the mean splicing efficiency of the host transcript. On the other hand, and in agreement with the analyzed distributions (Figure 2; Suppl. Fig. S2, S3), chromatin states of mRNA loci do not contribute to defining chromatin association of nascent mRNAs (Figure 4B). Similarly to lncRNAs, exon density and splicing efficiency of the worst spliced intron predict mRNA chromatin dissociation, while high SNRP70 enrichment over the transcription unit predicts slow-released mRNAs as well, suggesting that chromatin association of slow-released mRNAs could be at least partially achieved through persistent U1 snRNP binding to inefficiently processed transcripts.

**Figure 4.**
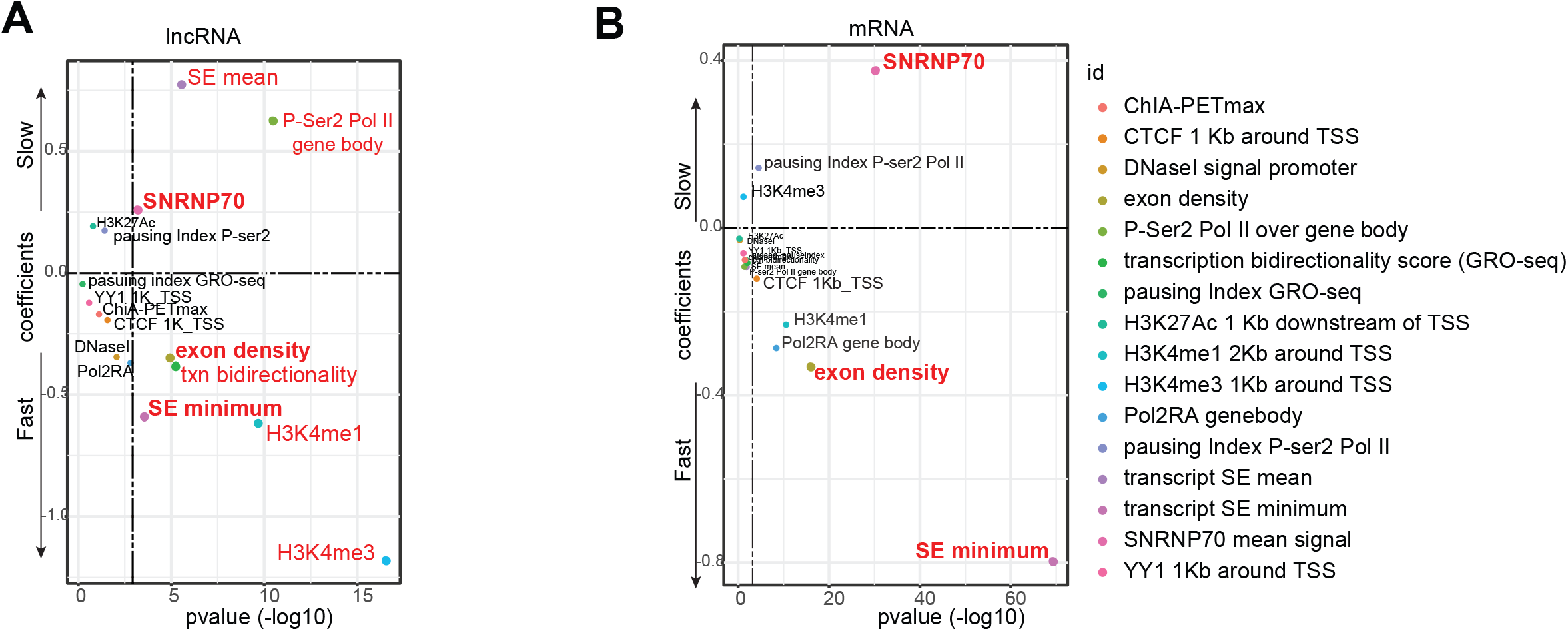
Contribution of distinct features to modeling chromatin (dis-)association of nascent RNA transcripts. (A) Logistic regression to predict fast vs. slow-released (chromatin-retained) lncRNAs. A 10x cross-validation was applied (best AUC 0.9275, average 0.8891). Coefficients bigger than 0.3 or smaller than −0.3 and with a p-value < 0.001 are marked red. (B) Same as in (A) but for mRNAs (best AUC 0.84, average 0.81).

Apart from logistic regression, we also applied linear regression to predict the chromatin association halftime (continuous value) as a multivariate function of several parameters (Supplementary Figure S4 A), as well as 2- class random forest (Supplementary Figure S4 B), reaching similar results regarding the weight of parameters in predicting fast versus slow-released transcripts.

### Distinct RNA binding proteins are predicted to bind transcripts of different degrees of chromatin association

We then asked whether lncRNAs of different degrees of chromatin association would interact with distinct RNA binding protein (RBP) activities. For this, we used the ENCODE-available eCLIP data^30^ from HepG2 cells as a proxy dataset. As most lncRNAs are expressed in a cell-type specific manner, we trained the *pysster* algorithm^31^ on mRNA or lncRNA sequences with overlapping RBP binding sites to acquire full-length transcript binding probabilities (by extracting the median score from positions that score above a pre-defined cutoff; Methods). We then incorporated these in random forest machine learning models to predict fast versus slow-released lncRNA or mRNA transcripts in 10 times cross-validation, with a mean accuracy of ~0.81 and ~0.8 respectively (Suppl. Figure S4C). Interestingly, we find RBPs with high binding probabilities which are commonly important in specifying chromatin association of both lncRNAs and mRNAs. These include factors with additional DNA binding activity (localizing to chromatin) like the KH-domain containing factors KHSRP and KHDRBS1, FUBP3 and SUGP2 which display increased binding probabilities for chromatin-retained/slow-released transcripts, either lncRNAs or mRNAs (Suppl. Figure S4 C lower panels). Interestingly, CSTF2 involved in 3’ end formation^32^ is also enriched in slow-released transcripts, perhaps reflecting persistent binding and unresolved RNA-protein complexes in the case of inefficient transcription termination and 3’ end formation. The exosome component EXOSC5 is also enriched in slow-released transcripts, implying chromatin-associated clearance of inefficiently processed nascent RNA transcripts. Among type-specific RBPs, DROSHA is an interesting candidate significantly enriched in fast-released lncRNAs, but not mRNAs, perhaps suggesting some involvement in promoting lncRNA chromatin dissociation in a causal manner (Suppl. Figure S4 C upper panels). Intriguingly, DROSHA was found important for pA-signal-independent transcription termination and 3’ end formation of lncRNAs serving as miRNA hosts^33^. Yet, the observed DROSHA enrichment (increased RNA binding probability) specifically in fast-released lncRNAs could also suggest post-transcriptional processing of nucleoplasmic-enriched lncRNAs. Although we do not find any significant enrichment of lncRNA miRNA hosts in the fast-released lncRNA category (since the numbers are quite small to infer statistical significance; only 34 of the 2,077 analyzed, expressed in MCF-7 lncRNAs host miRNAs), a more careful and closer examination would be required to conclude about microprocessor involvement in lncRNA transcription termination (and 3’ end formation) as an applying mechanism. Additional lncRNA-specific factors with increased RNA binding probabilities predictive for fast-released lncRNAs are NONO (involved in splicing), and XRN2 and CSTF2T, involved in transcription termination and 3’ end formation^34^. Since all three of them have DNA binding activity and localize to chromatin, this suggests that their predicted binding could be co-transcriptional and their activity may contribute to promoting chromatin dissociation of nascent lncRNA transcripts. Experimental examination by assessing the chromatin (dis-)association of nascent RNA transcripts in differential conditions upon RBP factor knock-down would validate these predictions and substantiate a specific candidate involvement in promoting efficient chromatin release or tethering.

## Discussion

lncRNAs constitute a large heterogeneous class with a broad range of functions in regulation of gene expression (regulation of transcription *in cis* and *in trans*), RNA processing and chromatin states^12,35^, while a common feature that distinguishes lncRNAs from mRNAs is reduced splicing efficiency^22^. The exerted functions of lncRNAs largely depend on their subcellular localization where they can differentially interact with distinct RNA-binding proteins and posit local target specificity. Previous computational efforts aimed to generate predictive models of lncRNA subcellular localization (nuclear versus cytoplasmic enrichment) using steady-state RNA-sequencing, and showed that inefficient splicing and intron retention is a major predictor of nuclear localization^24^. It is however an outstanding question what underlies the observed lncRNA chromatin enrichment (usually referred to as chromatin retention or chromatin tethering). lncRNAs may remain tethered to chromatin via ongoing Pol II transcription^10^ (since inhibiting Pol II transcription elongation abolished lncRNA chromatin tethering^10^), while the function of chromatin bound *cis* acting lncRNAs in regulation of proximal gene expression and local chromatin structure is mostly coupled to their ongoing transcription^11,36,37^. DNA elements in the *cis*-acting, chromatin-tethered lncRNAs may be key: for instance, loop interactions between the promoter of the chromatin-tethered lncRNA *PVT1* and its intragenic enhancers antagonize interactions with the neighboring *MYC* gene promoter^38^.

Yin et al. (2020)^14^ implicate persistent U1 snRNP binding as a means of lncRNA chromatin tethering, which relies on U1 site enrichment in lncRNA exons, depletion of 3’ splice sites and/or inefficient splicing, and U1 snRNP70 protein interactions with transcriptionally engaged Pol II. Interestingly, a previous study indicated that the overall lower splicing efficiency of lncRNAs (compared to mRNAs) is not due to defects in the U1-PAS axis which is very similar to mRNAs^22^. In agreement, we also find here SNRNP70 recruitment as a major predictive factor of lncRNA chromatin retention. In Yin et al. (2020), U1 inhibition dampened the chromatin association of both well and poorly spliced lncRNAs, suggesting that a kinetic effect due to delayed release of unspliced (or inefficiently/poorly spliced) nascent RNA cannot be the major determinant for lncRNA chromatin retention. In agreement, the transcript’s mean co-transcriptional splicing efficiency is a major predictor of chromatin dissociation for mRNAs but not lncRNAs. However, the splicing efficiency of the worst spliced intron per transcript has a significantly high coefficient in predicting chromatin dissociation, suggesting that it might function as a “bottleneck” for lncRNA chromatin release. Thus, nascent RNA splicing kinetics may at least partially contribute to lncRNA chromatin dissociation. Future experimental examination by point-mutating specific splice sites to enhance (or abolish) splicing will help to definitely validate the impact of co-transcriptional splicing kinetics on lncRNA chromatin release.

An important finding is that lncRNAs transcribed from active enhancers display increased degree of chromatin dissociation. This implies that the commonly termed “enhancer-associated” lncRNAs (or elncRNAs^7^, equivalent to la-EPCs^3^) do not remain chromatin associated. Instead, chromatin dissociation is an important feature which might underlie their function and impact enhancer activity. Again, it is noteworthy that single locus experimental validation aiming to alter the degree of lncRNA chromatin association will allow to examine the effect on cognate enhancer activity and target gene expression. So far, decrease in lncRNA chromatin tethering was achieved transcriptome-wide by inhibiting U1 snRNP^14^ (without examining the associated effects on putative *cis* targets), but it remains to be experimentally analyzed what is the effect of enforced elncRNA chromatin retention on cognate enhancer activity and target gene expression. This can be achieved either in modified cell lines by CRISPR gene editing or by targeting functional transcription termination and polyadenylation sites with blocking oligonucleotides for a short period of time to avoid secondary dampening effects on transcriptional activity. Modifying donor and acceptor splice sites of individual lncRNA loci by inserting point mutations should allow to experimentally validate the correlation between splicing efficiency and chromatin dissociation. It would also be highly relevant under such experimental conditions to examine alterations in local chromatin states and loop conformation, so as to characterize chromatin structure associated effects caused by enforced lncRNA chromatin tethering on enhancer functionality. Of great interest will be to draw conclusions on cognate enhancer activity after locus manipulation leading to altered locus-specific lncRNA chromatin-tethering without affecting splicing activity. This can be achieved for instance by interfering with 3’ end formation leading to increased transcriptional readthrough and suppressing nascent RNA transcript release^39^.

We note here that our approach to couple nascent RNA sequencing with chromatin fractionation at different pulse-chase time points would benefit by additionally applying long RNA sequencing of chromatin-associated and released nascent transcripts. The 3’ ends of long reads represent the position of Pol II at full-length (non-fragmented) synthesized nascent RNA transcripts, and this technique was recently employed to corroborate that co-transcriptional splicing greatly enhances mammalian gene expression (Reimer et al., biorxiv 2020). Previous experimental and computational studies focused at understanding nuclear retention of lncRNAs^24,40^. In these predictive models, inefficient splicing was a major factor contributing to lncRNA nuclear retention^24^. Here, by combining chromatin fractionation with sequencing of nascent RNA from the chromatin-associated and nucleoplasmic fraction at different pulse-chase time points and by employing machine learning we show that splicing of the least efficiently processed intron per transcript may act as a ‘bottleneck’ for efficient nascent RNA transcript chromatin release. Other factors like U1 snRNP (SNRNP70) binding, coupled with inefficient splicing, contribute to lncRNA chromatin retention as it was recently demonstrated in mESC^14^. We additionally show that lncRNAs transcribed from active enhancers do not remain chromatin tethered but rather display increased chromatin dissociation efficiency. Essentially elncRNAs are enriched in fast-released lncRNA transcripts, thus increased chromatin dissociation efficiency in addition to splicing^3,7^ may contribute to shaping enhancer activity and regulation of target gene expression. The latter may be accomplished upon chromatin dissociation of the nascent lncRNA transcript forming or affecting regulatory protein interactions targeting gene expression *in cis* within the spatial proximity of pre-established chromosomal loops.

---

**Supplementary Figure 1.**

(A) Nascent RNA sequencing read coverage over the last exon from all pulse-chase time points.

(B) Same as in (A) but after splitting in the 3 groups of fast, medium and slow-released transcripts.

(C) K-means clustering of all analyzed transcripts (n = 18,837) using the chromatin association ratios with k = 3 defines 3 clusters corresponding (from top to bottom) to ‘slow’, ‘fast’, and ‘medium’-released transcripts.

(D) Boxplot distribution of chromatin-association halftimes (calculated by fitting the exponential decay curve) for the k-means clustering-derived groups.

(E) Distribution of transcript length (fast vs. slow mRNAs p-value < 2.2e-16, fast vs. slow lncRNAs p-value 2.387e-12).

(F) Left panel: Distribution of number of exons per transcript (p-value 0.003545 fast vs. slow mRNAs, NS for lncRNAs). Right panel: Distribution of exon density (nr of exons per Kb) (p-value = 0.0009458 fast vs. slow lncRNAs, p-value < 2.2e-16 fast vs. slow mRNAs).

(G) Chromatin association halftime (extracted by fitting the chromatin association ratios on an exponential decay curve at p-value <0.05; Methods) for the three groups of fast, medium and slow released.

(H) Chromatin association ratios at steady state (log2 CHR/NP) for the 3 groups.

(I) Distribution of distances to closest antisense PCG TSS for the 3 groups of fast, medium and slow-released lncRNA transcripts.

(J) Cumulative distribution function curves of lncRNA interdistances to closest antisense PCG TSS for the 3 groups of fast, medium and slow-released lncRNA transcripts.

(K) Cumulative distribution function plots of intron splicing efficiencies from all time points and samples, at chromatin (left panel) and nucleoplasm (middle panel), and the respective boxplot distributions (right panel).

(L) Boxplot distribution of normalized post-transcriptional splicing efficiency at intron (left panel) and transcript level (middle and right panels; extracted as the mean normalized post-transcriptional splicing efficiency per transcript).

(M) UCSC screenshot from the GATA3-GATA3-AS1 locus.

**Supplementary Figure 2.**

(A) Transcript mean psi value (left) and median (right).

(B) Pausing index for lncRNAs and mRNAs (p-value < 2.2e-16) measured by extracting the ratio of P-Ser2 Pol II coverage 500 nt downstream of TSS to gene body.

(C) Same as in (B) but split for fast, medium and slow-released transcripts (fast vs. slow lncRNAs p-value 2.384e-08, NS for mRNAs).

(D) Pausing index using strand-specific GRO-seq read coverage (fast vs. slow released lncRNAs p-value 1.117e-15, NS for mRNAs).

(E) Strand-specific GRO-seq read coverage 1 Kb downstream of TSS (fast vs. slow lncRNA p-value = 6.185e-13, NS for mRNAs).

(F) Metagene analysis of GRO-seq strand-specific read coverage for the different groups of RNA transcripts, ±3 Kb around TSS.

(G) Transcription bidirectionality score extracted using GRO-seq (log2 antisense/sense read coverage 1 Kb around TSS; fast vs. slow lncRNA p-value < 2.2e-16, NS for mRNA).

(H) Distribution of interdistances of ContextMap extracted pA site to annotated transcript 3’ ends.

(I) Metagene analysis of average P-Ser2 Pol II density −500 bp to +5 Kb around TES of lncRNAs and mRNAs (left panel), and boxplot distribution of the corresponding transcription termination indices (extracted as the density ratio of 2.5 Kb downstream of TES to gene body (Methods); right panel).

(J) Metagene analysis of average GRO-seq read coverage profile (only the sense strand plotted) around the TES of grouped RNA transcripts; raw (upper panels), and after normalization of the average profile to value at nucleotide position zero (TES).

(K) Transcription termination index (NS)

(L) Travel index (fast vs. slow mRNAs p-value < 2.2e-16; fast vs. slow lncRNAs p-value 0.01248).

**Supplementary Figure 3.**

(A) Metagene profiles for different groups of transcripts (Id* label under panel C) of the average H3K4me1 to H3K4me3 ratio in a window ±3 Kb around TSS (left panel), and boxplot distribution of the overall H3K4me1 to H3K4me3 ratio 2 Kb downstream of TSS (right panel, p-value < 2.2e-16 fast vs. slow-released lncRNAs, p-value = 0.002769 fast vs. slow-released mRNAs).

(B) Average H3K9me3 (left) and H3K27me3 profiles around TSS of different groups of transcripts (Id* label under panel C).

(C) Average POL2RA profiles around TSS of different groups of transcripts.

(D) Average CTCF profiles around TSS of different groups of transcripts (first three panels), and boxplot distribution of CTCF enrichment ±1 Kb around TSS (fourth panel, p-value < 2.2e-16 fast vs. slow-released lncRNAs, p-value 6.032e-08 fast vs. slow released mRNAs).

(E) Average profiles of YY1 binding probability ±1 Kb around the TSS of different groups of transcripts (left panel, color Id* label under panel C), and the respective boxplot distributions of YY1 binding probability ±1 Kb around TSS (right panel, p-value < 2.2e-16 fast vs. slow-released lncRNAs, NS for mRNAs).

(F) Boxplot distributions of promoter ChIA-PET score, extracted as the sum of scores of the ChIA-PET nodes overlapping the promoter (±2 Kb TSS) (p-value 2.375e-09 fast vs. slow released mRNAs and 3.039e-11 fast vs. slow-released lncRNAs).

**Supplementary Figure 4.**

(A) Linear regression models (lm) run with 10 x cross-validation to predict chromatin association halftime (as a continuous value) of lncRNAs (left panels) and mRNAs (right panels) by incorporating several parameters (significant parameters with a coefficient p-value < 0.001 are red-marked).

(B) Two-class random forest run with 10 x cross-validation to predict fast vs. slow released lncRNAs (upper panels, best model accuracy 0.91, mean accuracy 0.86) and mRNAs (lower panels, best model accuracy 0.81, mean accuracy 0.77).

(C) Two-class random forest run with 10 x cross-validation to predict fast vs. slow released lncRNAs (upper panels, best model accuracy 0.808, mean accuracy 0.769) and mRNAs (lower panels, best model accuracy 0.795, mean accuracy 0.776) by incorporating 100 RBP whole transcript binding probabilities (*pysster* predictions). Mean Decrease Accuracy and Mean Decrease Gini values of the top best 30 factors are shown.

(D) Boxplot distribution of whole transcript binding probabilities (*pysster* predictions) for factors important either for predicting fast vs. slow-released lncRNAs (upper panels), fast vs. slow-released mRNAs (middle panels) or both types (lower panels). Student’s *t.test* p-values are noted (red for fast vs. slow lncRNAs and blue for fast vs. slow-released mRNAs).

## Acknowledgements

We thank Rutger Gjaltema and Edda Schulz for helpful discussions and comments on the manuscript. This work was supported by the DFG Grant MA 4454/3-1 to A.M.

## AUTHOR CONTRIBUTIONS

N. and A.M. conceived and planned the study with input from UAVØ. E.N. designed the computational and experimental pipelines, performed experiments and computational analyses. S.B. implemented the *pysster* method and contributed to data analyses. A.M. supervised computational analyses. E.N. and A.M. wrote the manuscript.

## Methods

### KEY RESOURCES TABLE

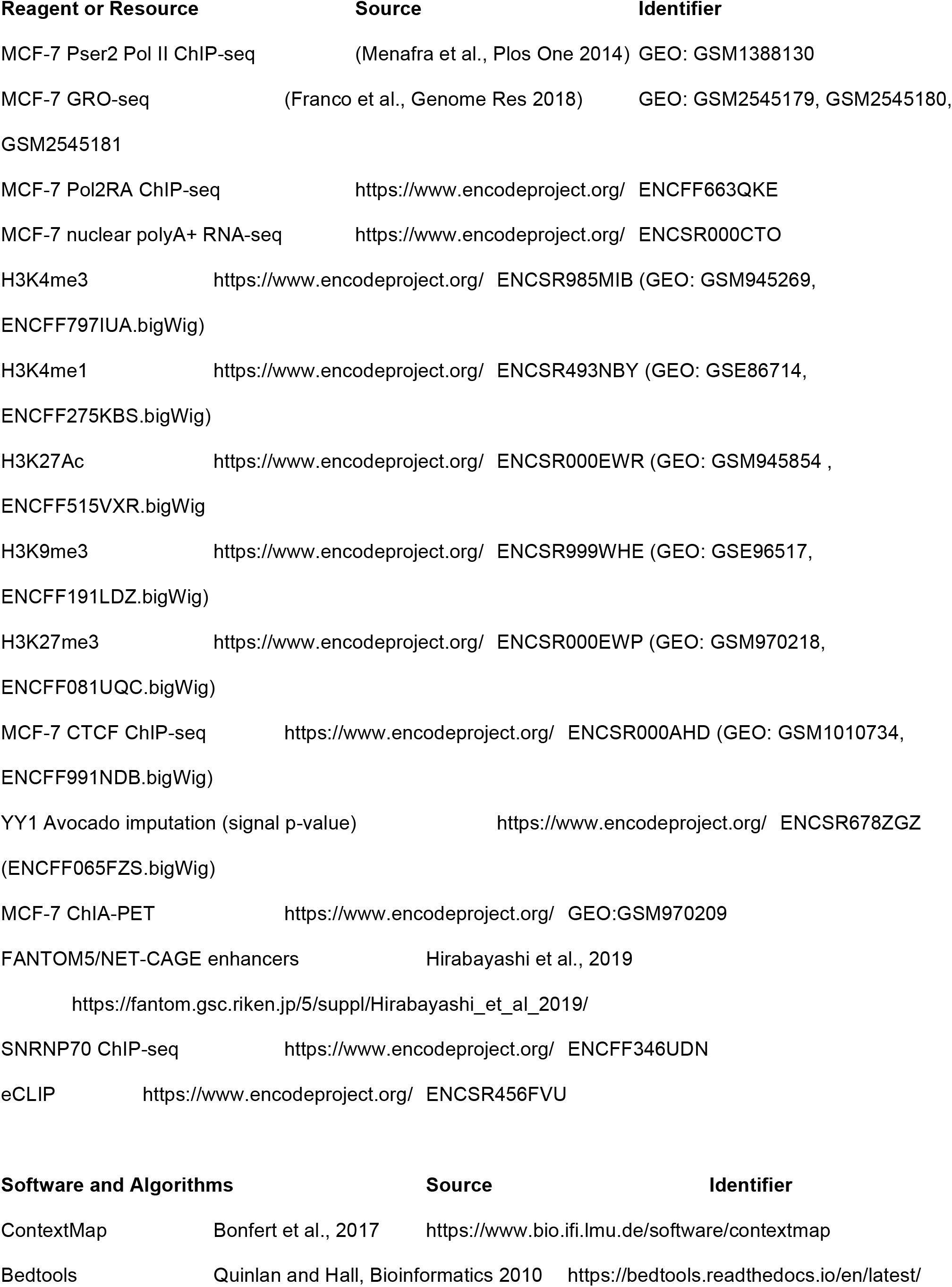
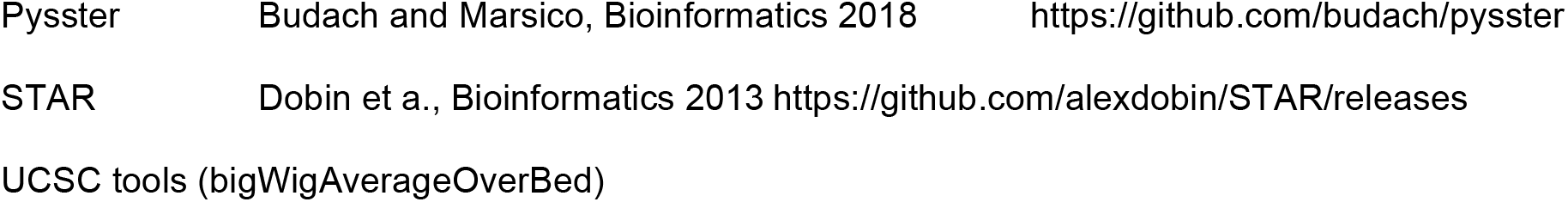

## METHOD DETAILS

### Extraction of transcript 3’ end site (TES)

We ran ContextMap v2.7.9 on paired-end MCF-7 nuclear polyA+ data (ENCODE) using Bowtie2 aligner and Bowtie2-build-l indexer, with parameters -mismatches 3 -seed 30 -maxhits 10 --polyA -t 8 -Xms4000M - Xmx30000M. This generated 39,991 ContextMap scored polyA sites. Nearby polyA sites were clustered with bedtools cluster –s –d 10, keeping the one with maximum score. Annotated transcript 3’ ends were assigned a ContextMap polyA site by fetching the closest with bedtools closest –s.

### Enhancer-associated lncRNAs in MCF-7

From the FANTOM5/NET-CAGE enhancers (n = 85,786) we extracted the ones that show evident bidirectional transcription in MCF-7 using GRO-seq (GSE96859) (bigWigCoverageOverBed mean0 coverage > 0.1 for both strands), resulting in 10,008 bidirectional actively transcribed enhancers. We then fetched closest transcript start site (TSS) to enhancer midpoints using bedtools closest –s and defined lncRNAs with an interdistance < 2000 bp as elncRNAs (n = 248 out of the 2077 analyzed).

### SNRNP70 occupancy over transcription units

As a proxy we used SNRNP70 ChIP-seq from HepG2 and by intersecting the intervals corresponding to full-length transcripts with ChIP-seq narrow peaks (ENCFF346UDN) we extracted a mean binding score per transcription unit.

### Nascent RNA sequencing combined with pulse-chase and chromatin fractionation

MCF-7 cells were seeded in P10 (6 plates per time point) and grown to ~80% confluency in 5% FCS, then labeled for 8 min with 1mM 4-thio-Uridine (4-SU). Cells were either immediately harvested (lifted intact in ice-cold PBS) or washed twice in PBS and chase was applied for 5, 10, 15, 20 min in 10 mM uridine diluted in growth medium. Chromatin fractionation was performed as in ref^9^. Briefly cells were lysed in 400 ul lysis buffer 0.15% NP-40 and lysate was loaded on 800 ul sucrose buffer for brief centrifugation. Pelleted nuclei were washed in ice-cold PBS, resuspended in 200 ul glycerol buffer and lysed in 0.6 M urea to fractionate chromatin from the nucleoplasmic fraction. RNA from the chromatin and nucleoplasmic fraction was extracted with acidic phenol (pH 4.5) and acidic phenol/chloroform. 3 ug of RNA were fragmented with 0.15 M NaOH final concentration for 25 min on ice. Prior the RNA fragmentation, 0.15 ng of the 4-SU-labeled and unlabeled spike-ins mix (as in the TT-seq protocol^15^) had been added to the 3 ug of RNA. The fragmentation reaction was stopped in 10 mM Tris pH 7.4, purified with RNeasy MinElute Spin columns and eluted in 45 ul TE buffer (Tris 10 mM pH 7.4, 1mM EDTA). 5 ul Biotin-HPDP/DMF 1 mg/ml were added (i.e. final concentration 0.1 mg/ml) and incubated for 2 hours at room temperature. Further steps of RNA purification, binding to T1 Dynabeads, washing and elution were done according to the A. Regev protocol^41^ (using 5 ug T1 Dynabeads for 2 ug 4-SU-biotinylated RNA), leading to library construction for Illumina sequencing.

### Mapping and spike-ins normalization

Reads were mapped to GRCh38 (gencode.v23.primary_assembly.annotation) and to ERCC92 sequences using STAR 2.5.4a with standard parameters. Only reads mapped to a single genomic location were kept (score 255). Three labeled (ERCC00043, ERCC00092, ERCC00136) and three unlabeled spike-ins (ERCC00002, ERCC00145, ERCC00170) had been added to each RNA sample. For each sample a ‘sizeFactor’ was extracted for spike-ins normalization as follows: each of the three labeled spike-ins read counts were normalized to the sum of the respective spike-in counts across the ten labeled samples (CHR 0, 5, 10, 15, 20 min and NP 0, 5, 10, 15, 20 min), and then the median value from the three normalized labeled spike-ins was extracted per sample (‘smoothened median’ = sizeFactor). For each labeled sample the cross-contamination value ‘epsilon’ was calculated as the sum of unlabeled spike-in read counts (U) to the sum of U plus the sum of labeled spike-in read counts (L): epsilon = cross-contamination = U/ (L+U). Strand-specific read counts over features were normalized to sizeFactor and feature length and multiplied by (1-epsilon). Fitted_counts = measured_counts/ sizeFactor(labeled_sample)/ feature_length * (1-epsilon), or Fitted_counts = measured_counts/ sizeFactor(labeled_sample)/ feature_length * L/ (L+U).

### Transcript dataset

We used GENCODE V29 lncRNA annotation (n = 8,992) supplemented with novel (non-overlapping GENCODE V29 lncRNAs annotation) lncRNA transcripts from *de novo* transcript assembly (n = 10,606) on chromatin-associated RNA-seq in MCF-7 (described in ref^9^; those are lacking protein-coding potential, are not overlapping protein coding genes, and have at least 1 splice junction). From this initial set we kept 3,671 lncRNAs with non-zero read coverage in all 12 sequenced samples. We also used 15,166 mRNA transcripts with non-zero read coverage in all 12 samples.

### Modeling chromatin dissociation

Strand-specific read counts over the last exon of the 18,837 transcripts were normalized to spike-ins and feature length (as described in the Methods section ‘Mapping and spike-ins normalization’). For each pulse-chase time point we extracted a ratio of chromatin (CHR) to chromatin plus nucleoplasmic (NP) normalized read coverage (CHR/ (CHR+NP)). We fit those ratios on an exponential decay using R function lm (log (x) ~ time), for timepoints [0,8, 13, 18, 23, 28] (ratio set to 1 at timepoint 0), which returns *intercept*, *k* and *p-value* of exponential decay fit. We kept 12,391 entries that fit the curve with a p-value <0.05 (of which 2077 lncRNAs, and 10,314 mRNAs). We defined a ‘chromatin association halftime’ as -(*intercept* + log (2)) / *k*. Based on the halftime values, we split the dataset in three equal-size quantiles corresponding to ‘fast’, ‘medium’ and ‘slow’ released nascent RNA transcripts.

### Splicing efficiency, SED and degree of post-transcriptional splicing

We measured intron splicing efficiency (SE or *thita* value) as in ref^20^ by extracting the ratio of split to split plus non-split reads overlapping 3’ splice sites of introns with at least one split and one non-spit read at the 3’ splice site (n = 154,467 high-confidence introns). We measured alternative splicing as in ref^21^ by extracting the ratio (*psi* value) of alternative split to constitutive split reads covering the high-confidence introns. We extracted co- and post-transcriptional splicing efficiency dynamics (SED) as in ref^21^, by subtracting the difference of splicing efficiency at 20 min pulse-chase from the splicing efficiency at 0 min and normalizing this to the splicing efficiency at 0 min [*SED = (SE_20min + 0.001 – SE_0min) / (SE_0min + 0.001)*]. We extracted the extent of post-transcriptional splicing relative to co-transcriptional as the difference of chromatin-associated splicing efficiency from the nucleoplasmic splicing efficiency, normalized to chromatin. This was done at intron and transcript level (mean value of the transcript’s high-confidence introns).

### Transcriptional indices (TSS-proximal pausing index and termination index)

We assessed transcriptional pausing index by extracting the ratio of strand-specific GRO-seq read coverage or P-Ser2 Pol II ChIP-seq density in the window 500 nt downstream of TSS to the gene body. Gene body was defined as the middle 50% of the interval TSS+500 to TES, as in ref^20^. Transcription termination index was measured as in in ref^20^ by extracting the length-normalized ratio of strand-specific GRO-seq read coverage (or Pol II ChIP-seq read density) in the window 2.5 Kb downstream of TES to gene body. Travel index was extracted as the ratio of read coverage in the interval [2.5 to 5 Kb] downstream of TES to the first 2.5 Kb downstream of TES.

### Machine learning models

Logistic regression to predict fast versus slow-released nascent RNA transcripts was ran on standardized parameters (R function *stdize()* of the package ‘*pls*’) using the R function *glm()* and ten-times cross-validation. Linear regression to model chromatin-association halftime as a continuous value was ran on standardized parameters using R function *lm()* and ten-times cross-validation. Random forest to predict fast versus slow-released nascent RNA transcripts was ran with R function *randomForest()* and ten-cross-validation, setting number of trees 1000 (*ntree* = 1000) and dataset-specific best *mtry* parameter. Best *mtry* was found using the function train() of the package ‘caret’times with a grid-search and ten-times cross-validation.

### RBP predictions (build *pysster* models and prediction scan summary)

To train *pysster* models we used ENCODE available eCLIP data from HepG2 cell line for 100 RNA-binding proteins (2 biological replicates). eCLIP peaks found in both biological replicates and with a log-fold enrichment > 2 over the input control were selected (5’ end of peaks are used as binding sites from now on). *Pysster* was used to train a multi-class convolutional neural network (CNN) classifier. We trained one model for each RBP, and each model was trained on 3 classes:

- class 1: sequences of length 400 centered at a binding site of the protein of interest
- class 2: randomly sampled sequences of length 400 from lncRNAs (lncRNA models) or mRNAs (mRNA models) that contain at least one binding site of the protein of interest (sequences were sampled such that they don’t overlap with class 1 though)
- class 3: sequences of length 400 centered at randomly selected binding sites of all other proteins to reduce the impact of eCLIP bias signal (no overlap with class 1 again)

In addition to the sequences itself, the CNNs also use the following additional data as input: (1) is sequence position 200 located in an exon or intron? (zero/one encoded), (2) distance of sequence position 200 to the TSS/TTS (normalized to the transcript length such that zero indicates overlap with the TTS and one overlap with the TSS). For each model a hyperparameter grid search was performed: 3 convolutional layers, kernels of length 12, 18 or 24 and 150 or 300 kernels per layer (all other *pysster* parameters were left at their defaults). A trained RBP model could then be applied to a transcript of interest as follows: using a sliding window approach (window size 400, step size 1) the score of belonging to class 1 was predicted for all bases of a transcript. All predictions > 0.66 were selected and their median was computed.

