## Supplementary figures and images for "Predictive modeling of long non-coding RNA chromatin (dis-)association"

### Supplemental Figure S1

#### Supplemental Figure S1

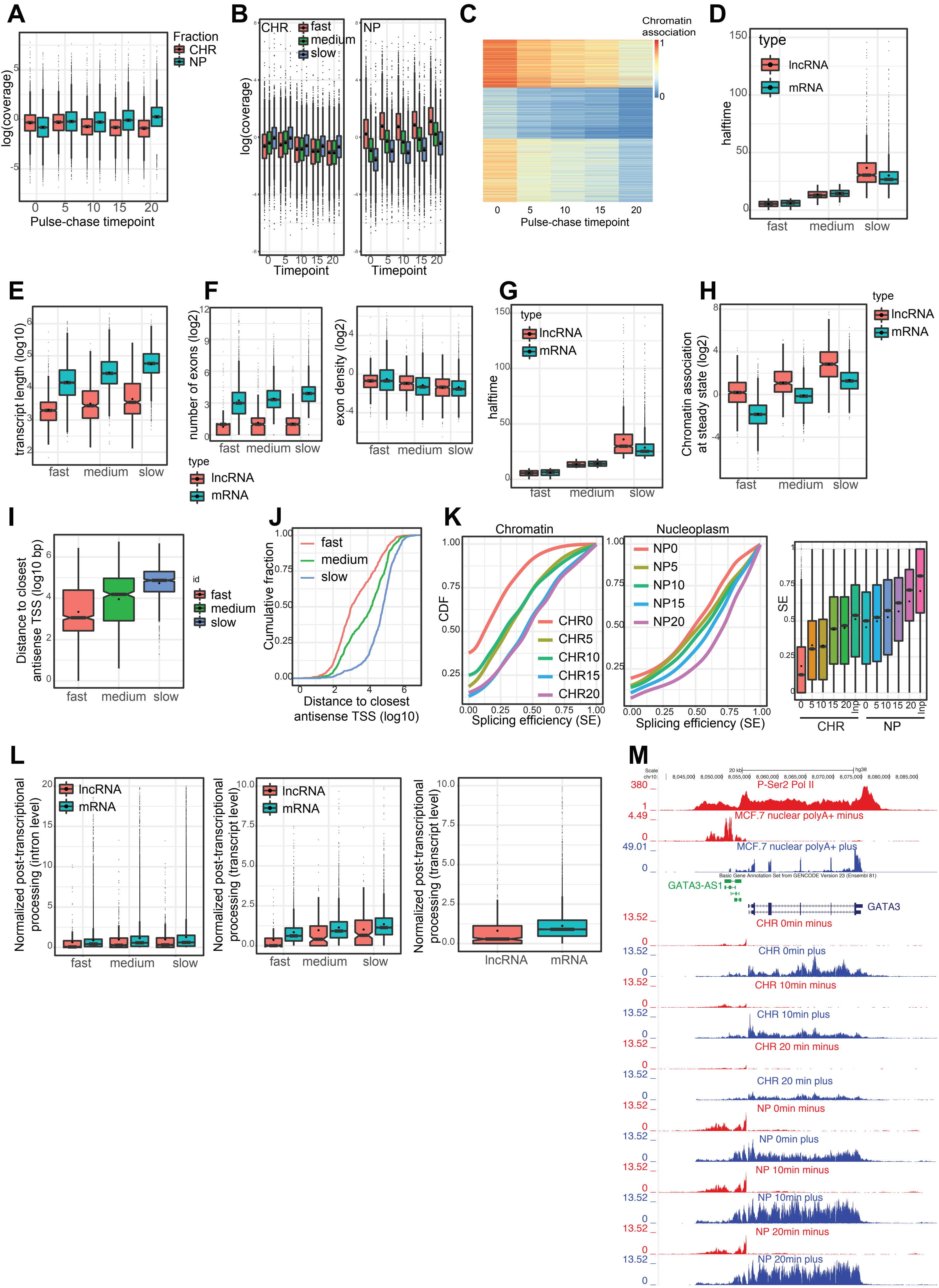

### Supplemental Figure S2

#### Supplemental Figure S2

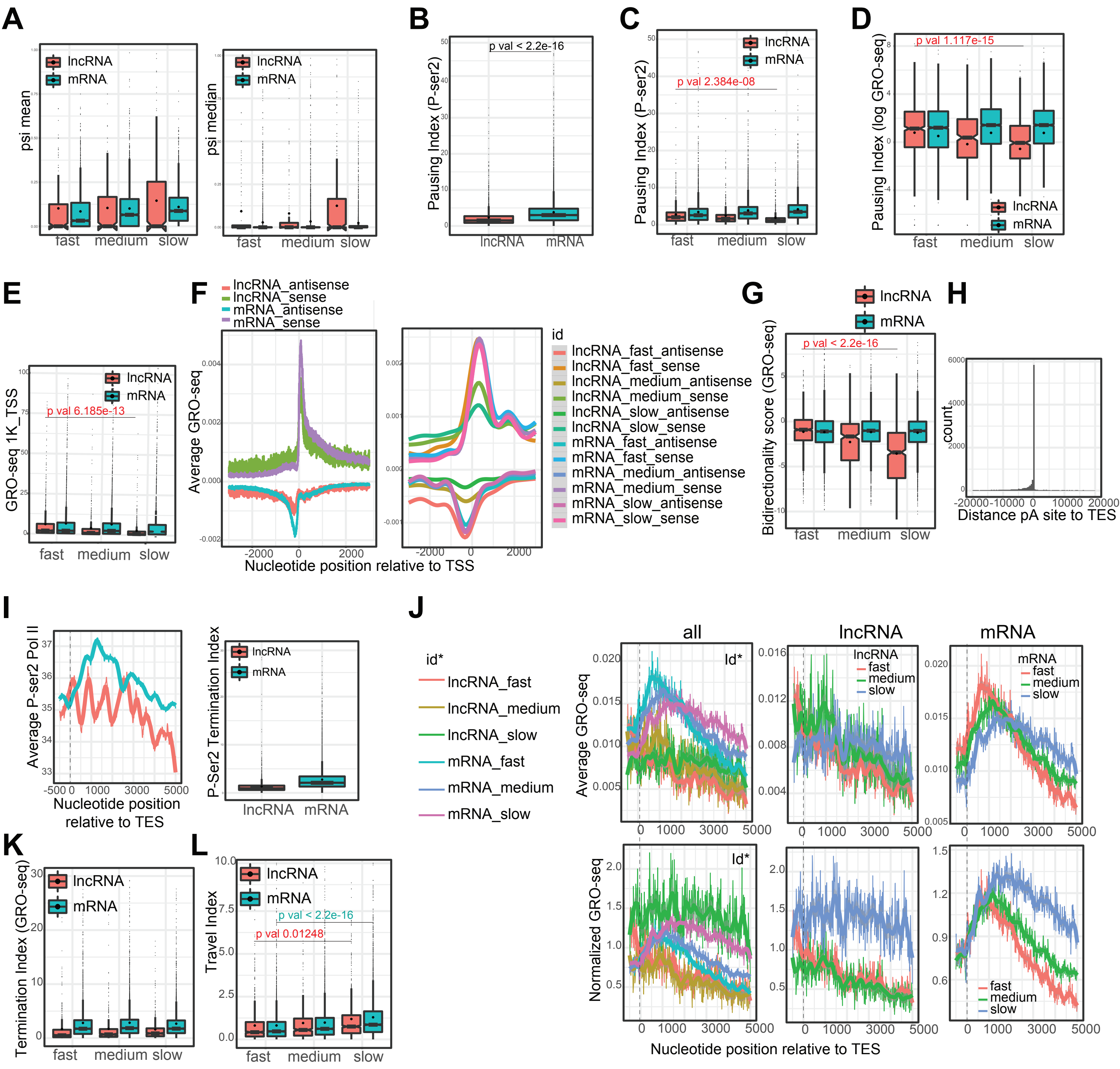

### Supplemental Figure S3

#### Supplemental Figure S3

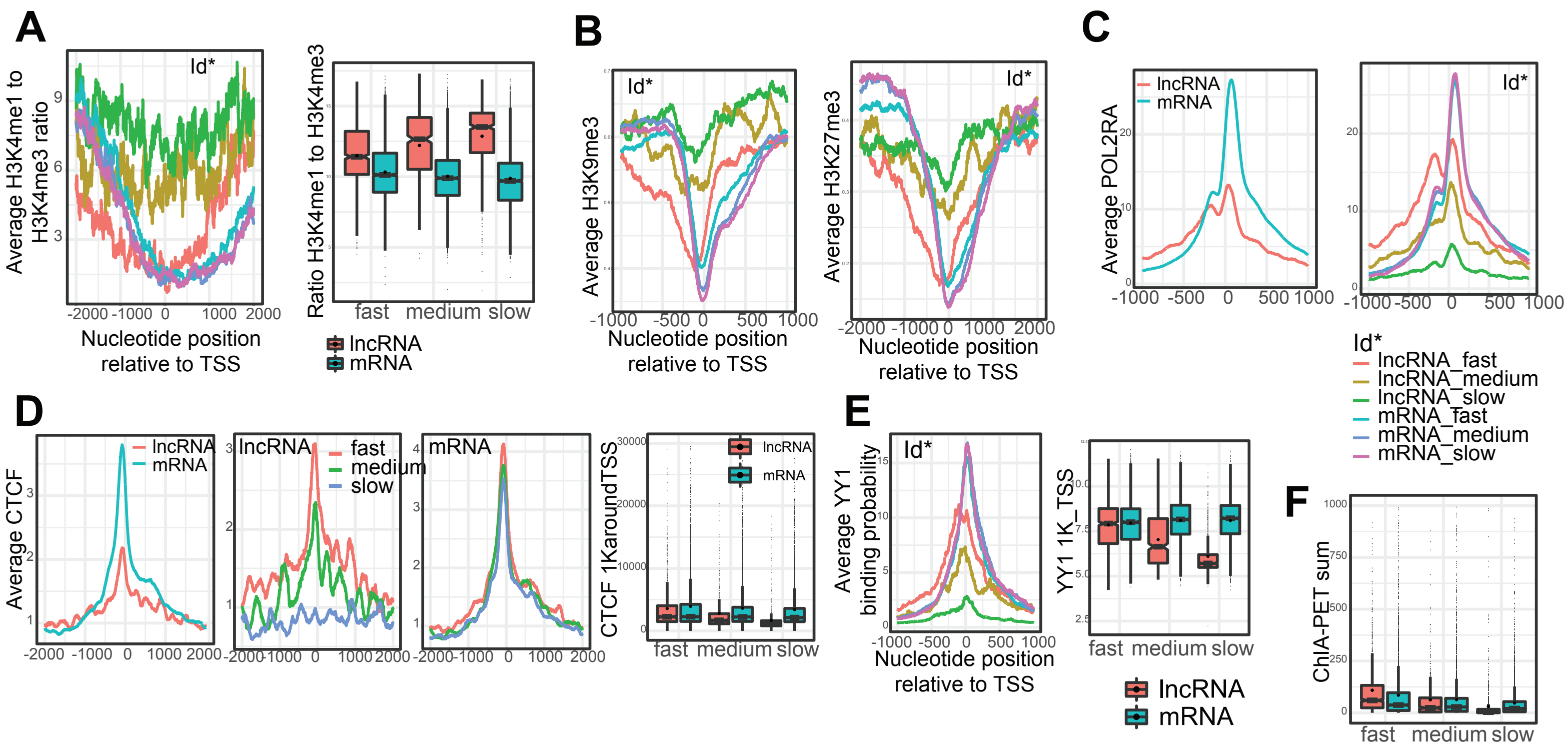

### Supplemental Figure S4

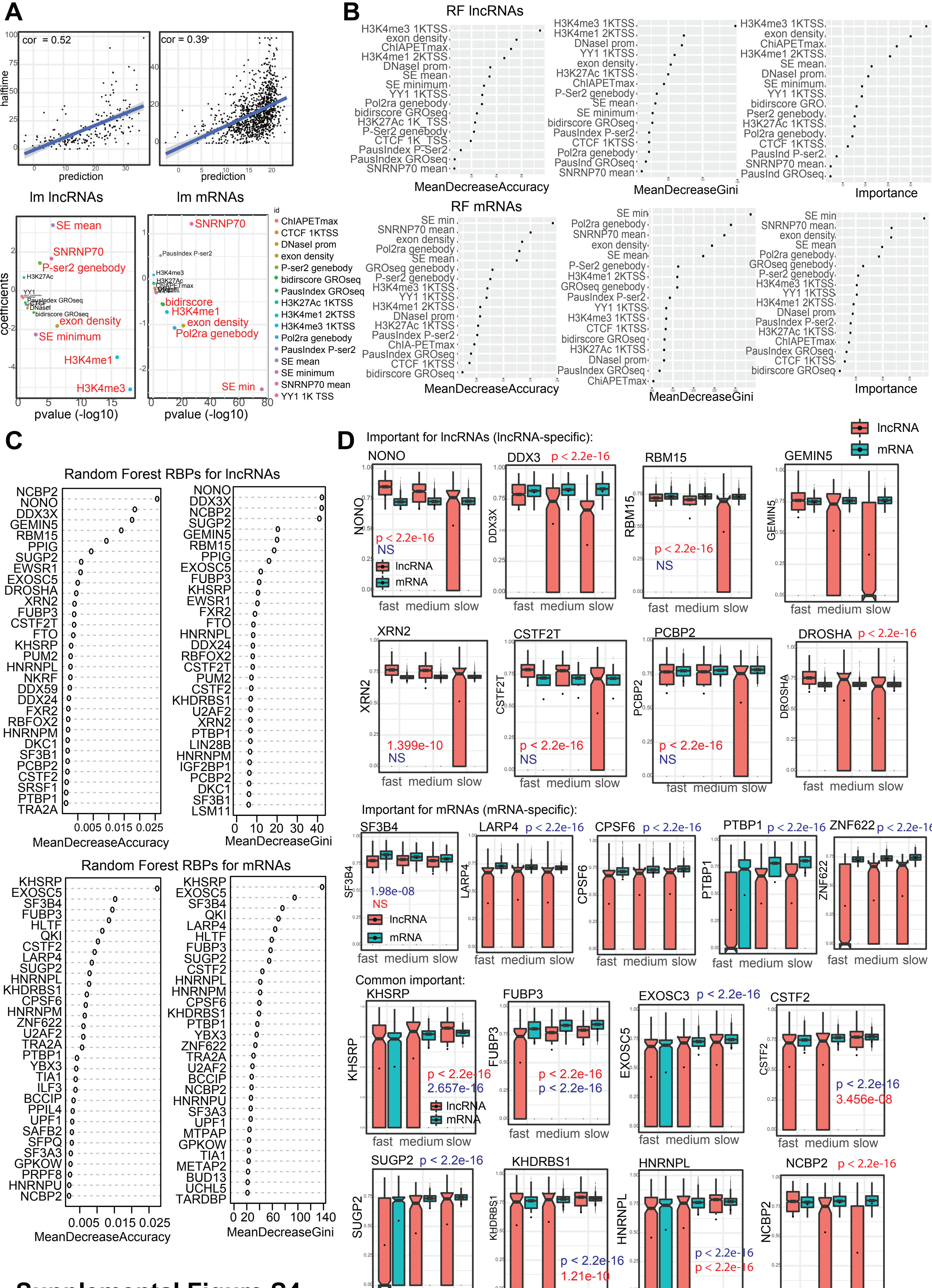


**Supplemental Figure S4**
